# Gene-environment interactions in face categorisation: experience with a nanny and oxytocin receptor genotype interact to reduce face categorization reaction times

**DOI:** 10.1101/2020.12.17.423340

**Authors:** Michelle Jin Yee Neoh, Peipei Setoh, Andrea Bizzego, Moses Tandiono, Jia Nee Foo, Gianluca Esposito

**Affiliations:** Psychology Program, School of Social Sciences, Nanyang Technological University, Singapore, Singapore; Department of Psychology and Cognitive Science, University of Trento, Rovereto, Italy; Lee Kong Chian School of Medicine, Nanyang Technological University, Singapore, Singapore; Human Genetics, Genome Institute of Singapore, Singapore, Singapore

**Keywords:** face categorisation, perceptual expertise, multiracial, oxytocin receptor gene, gene-environment interaction

## Abstract

Human faces are relevant stimuli that capture attention, provide information about group belonging and elicit automatic prepared responses. While early experiences with other race faces plays a critical role in acquiring face expertise, the exact mechanism through which it exerts its influence is still to be elucidated. In particular, the influence of genetic factors and the role of a multi-ethnic context has not been explored. The aim of this study was to investigate how caregiving experiences with nannies and oxytocin receptor gene (OXTR) genotypes interact in regulating other-race categorisation mechanisms in adults. Information about single nucleotide polymorphisms of the OXTR (rs53576) and experiences with own- and other-race nannies was collected from 89 Singaporean adults, who completed a visual categorization task of face stimuli (Chinese or Javanese). Participants were grouped into A/A homozygotes and G-carriers and assigned a score to account for the type of nanny experience. A General Linear Model was used to estimate the effect of nanny experience, genetic group and their interaction on categorization reaction time. A significant main effect of the nanny experience (p<.001) and of the interaction between genetic group and experience (p=.008) was found. Post-hoc analysis revealed a significant negative correlation between nanny experience and reaction time for A/A homozygotes (r=−0.52, p<.001) but no significant correlation for G-carriers. In summary, a significant gene-environment interaction on face categorization was found. This finding appears to represent an indirect pathway through which genes and experiences interact to shape mature social sensitivity in human adults.

**Highlights:** Early nanny experience interacts with oxytocin receptor genotype in affecting the speed of face categorisation.

Individuals with other-race nanny experience show faster categorisation response times. Gene-environment interactions are present in face categorisation.

## 1. Introduction

Faces have a homogenous positioning of internal elements regardless of the individual, and early in life, human infants are attracted to faces and face-like stimuli (Goren et al. 1975, Morton & Johnson, 1991). From an evolutionary perspective, Seligman's theory of ‘preparedness’ proposes that stimuli which are critical for survival, such as threat stimuli, elicit automatic prepared responses (Ohman & Mineka, 2001). Extension of this theory predicts that the human brain would show innate predispositions to react not only to threat stimuli but to all biologically salient stimuli independent of their valence (Scherer, 2001). In this view, human faces represent highly biologically relevant stimuli that capture attention, provide information about group belonging and to which humans might be prepared to respond. Humans are thus equipped with a rapid and automatic process to categorize faces as belonging to one’s own social group or not (Pascalis, Loevenbruck, Quinn, Kandel, Tanaka & Lee, 2014).

People categorise others based on social categories – namely, age, race and sex (Zhao & Bentin, 2008) using particular cues. For example, face categorisation decisions regarding race are influenced by skin colour (see Maddox, 2004 for review) and other facial features are used (e.g Dunham, Stephanova, Dotsch & Todorov, 2014; Stepanova & Strube, 2012). The mechanisms underlying social categorisation are complex. The accuracy of social categorisation is built upon the ability to learn the perceptual features which best indicate group membership.

While experience plays a critical role in acquiring face expertise, the mechanisms through which it exerts its influence is still to be elucidated. An individual’s ability to categorize faces based on social concepts such as race stems from early, repeated experiences and develops through socialization processes (Dunham, Chen & Banaji, 2013; Rutland, Cameron, Milne & McGeorge, 2005; Xiao et al, 2015). Social categorization perspective centers around the premise that differential processing is employed for ingroup vs outgroup members (Sporer, 2001). When a face is encountered, rapid categorization of the face as belonging to an ingroup or outgroup occurs (Levin, 2000). Ingroup faces tend to be processed at an individual level, whereas outgroup faces tend to be processed at a more superficial, categorical level (see review by Sporer, 2001). In addition, differential allocation of attention to outgroup members has been suggested to lead to weaker encoding of features relating to outgroup members in comparison to ingroup members (Rodin, 1987).

The categorisation-individuation model proposes how categorisation and experience interact in explaining differences in face encoding processes and differences in memory for ingroup vs outgroup faces (Hugenberg et al, 2010). The model proposed the role of (i) perceiver characteristics, specifically familiarity and experience with individuals from the race, and (ii) social categories, specifically differential attention to category-diagnostic facial characteristics in facilitating categorization and individuation processes. The model suggests that facial features differentiating between categories are first attended to, and experience with individuating a particular category facilitates selective attention to features which best distinguish between members of that category. However, the categorisation-individuation model is limited in that it does not explore biological factors such as genetics or neural processes underlying face recognition.

### 1.1 Role of experience in face categorisation

Real-life experiences have been proposed to play a critical role in acquiring face expertise (Lee, Anzures, Quinn, Pascalis & Slater, 2011). Greater categorisation accuracy is achieved with experience interacting with members of a particular category, possibly due to increased opportunities to learn relevant facial features for distinguishing group boundaries. Nine-month-old White infants could categorise and individuate White faces but not Chinese faces, indicating the role of experience in categorisation of face race (Anzures et al, 2010). Furthermore, there has been some preliminary evidence indicating the role of early caregiving experience plays in face processing. Tham, Woo & Bremner (2018) found that three- and four-month-old infants showed better recognition for faces of the same race and gender of their primary caregiver. In addition, the study found additional evidence indicating that infants with female other-race nannies showed better recognition for nanny-race faces compared to those without, suggesting the role of caregiving experience and other-race contact on face processing.

### 1.2 Role of genetics on face categorisation

Beside the effect of experience on face processing abilities, a significant body of research documents the importance of some specific genetic factors that are associated with social cognition and social sensitivities to the environment (see Skuse & Gallagher, 2011). In line with principles of the categorisation-individuation model regarding differences in face processing, we propose that genetic factors which have been associated with social cognitive processes could influence race-based face categorisation. The oxytocin system has been implicated in human socially related personality traits and behaviors (Donaldson & Young, 2008; Meyer-Lindenberg, Domes, Kirsch & Heinrichs, 2011). Specifically, the oxytocin receptor gene (OXTR) has been identified as a candidate gene regulating attachment-related behaviors and social cognitions in humans where the link between single nucleotide polymorphisms (SNPs) in this gene and sociality has been examined Ebstein, Israel, Chew, Zhong, Knafo, 2010; Gillath, Shaver, Baek & Chun, 2008). One such SNP is rs53576 (G/A), which previous studies have found G allele homozygotes (GG) to be associated with higher (i) self-reported empathy (Rodrigues, Saslow, Garcia, John & Keltner, 2009), (ii) prosocial features (Tost et al, 2010) and (iii) general sociality as rated by peers (Kogan, Saslow, Impett, Oveis, Keltner & Saturn, 2011) as compared to A allele carriers (A/A or A/G). While these findings suggest the influence of the OXTR genotype on social behaviors and cognition, the effect of this SNP genotype specifically on race-based face categorization has not been studied previously. Hence, this study investigated the role of the rs53576 in the OXTR gene on race-based face categorisation and its possible interactions with early caregiving experience.

### 1.3 Exposure to faces in a multiracial context

Majority of the studies on face processing have been conducted in predominantly monoracial populations living in racially homogeneous environments who generally lack experience with other-race faces (Valentine & Endo, 1992; Levin, 1996; Zhao & Bentin, 2008, Feng et al, 2011) and there are fewer studies looking at participants from multiracial societies. Hence, there appears to be a gap in research examining face processing in populations with greater experience with faces of different categories – specifically, race – especially experience that occurs early on in life. In this respect, research conducted in multiracial societies where individuals have consistent, early exposure to a diverse range of faces of different races can help to address this research gap. In line with the discussion on exposure and experience with faces of other categories, it is likely that results on face processing will show different patterns. For example, Woo et al (2020) found that both Malaysian Chinese children and adults did not exhibit an other-race categorization advantage as they categorized own-race Chinese faces more rapidly than high-frequency Malay faces (i.e. the predominant racial group in Malaysia). Therefore, the present study conducted in Singapore, a multiracial society, (resident population consists of 76% Chinese; 15% Malay; 7.5% Indian; and 1.5% other races; Strategy Group, 2019) aims to examine early caregiving experience on race-based face categorisation in Singaporean adults. In this study, early caregiving experience is defined as being cared for during infancy or childhood by a kin or non-kin nanny other than one’s own parents, which is a common practice in the country.

### 1.4 Present study

The present study aims to explore how genetic factors, childhood exposure to stranger and other-race faces and their interactions influence the mechanisms involved in race-based face categorisation, focusing on a particular multi-racial context. Specifically, we investigated the effects of early caregiving experiences and the genotypes of OXTR in Singaporean adults in influencing adults’ categorization of own-race and other-race faces (see Figure 1).

**Figure 1.**
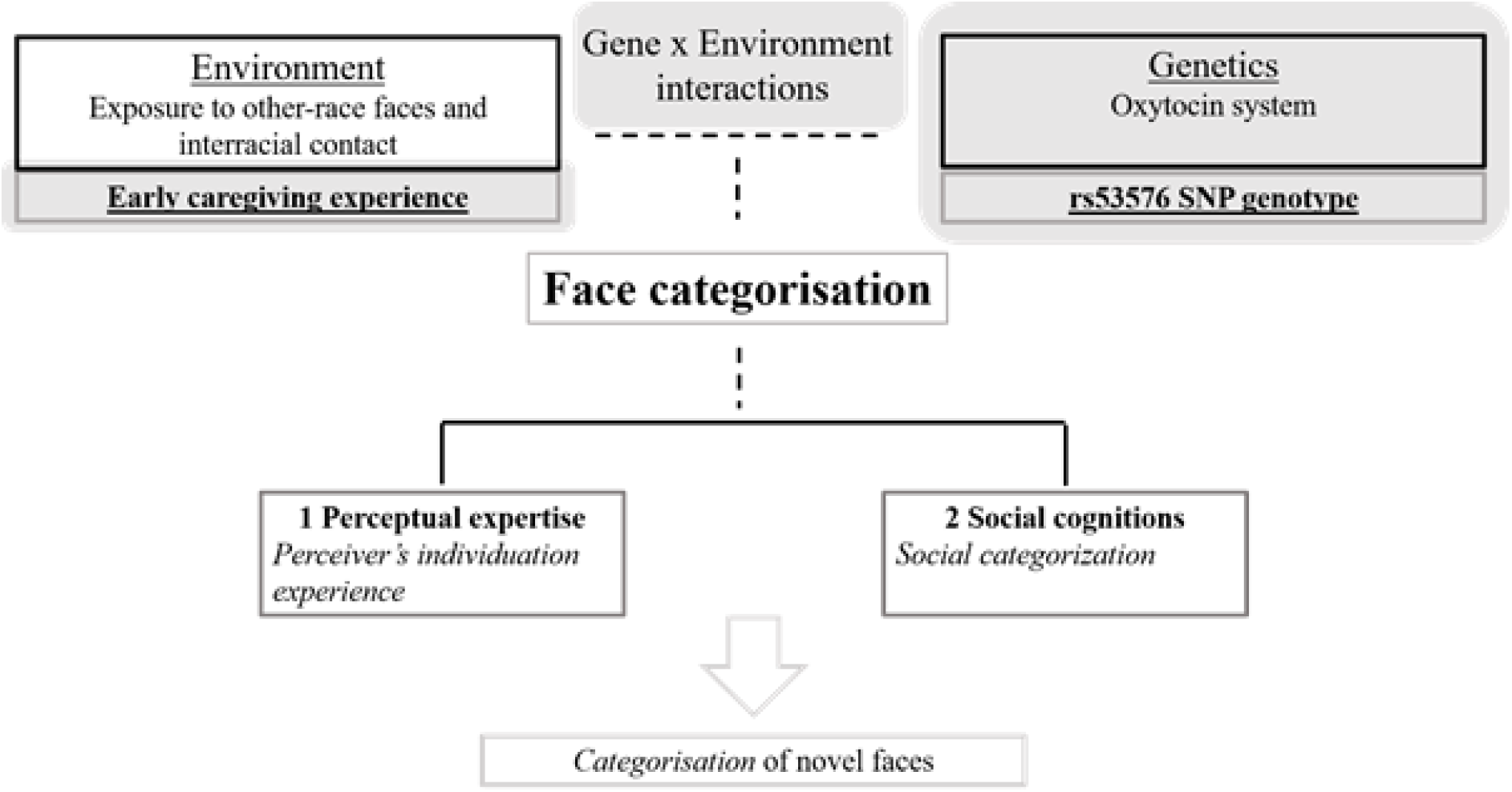
Diagram of existing theoretical frameworks of face categorisation with proposed additions and key research variables of the present study highlighted in grey in the figure.

Regarding the environmental factor, we focused on early caregiving experiences with same- or other-race nannies or no caregiving experience.

Regarding the genetic factor, the present study examined the oxytocin system, which has been implicated in human socially related personality traits and behaviours. Specifically, the rs53576 SNP genotype was examined as the G allele of rs53576 appears to be associated with prosocial traits including sociality and social cognition (Tost et al, 2010; Kogan et al, 2011; Li et al, 2015).

The time taken for visual categorization of the face stimuli by race was measured to determine the speed of processing in categorising faces by race. Due to the multiracial nature and population make-up of the Singaporean context, participants are likely to have a high exposure to faces of different races. As a result, the other-race categorisation advantage is likely to be reduced in Singaporean participants. A preliminary analysis is first conducted to check for significant differences in categorisation reaction times for own and other race faces. Firstly, we hypothesized that there will be significant differences in the reaction times for race categorization between individuals with different exposure to stranger or other-race faces. In line with the perceptual expertise hypothesis and findings from Woo et al (2020) of overall faster categorisation response times in Chinese participants, we expect that individuals with other-race nanny experience will show faster categorization reaction times compared to those with own-race nanny or no experience. Then, we hypothesized that OXTr genotype will interact with early caregiving experience in influencing categorisation responses. In line with findings of the association between G allele and prosociality, we expect that G allele carriers will have greater social interactions and contact with others. As such, early caregiving experience with nannies constitutes a more significant source of interracial contact for A/A carriers compared to G/ carriers. Hence, faster categorisation reaction times for individuals with caregiving experience with other-race nanny compared to those with own-race nanny or no experience will only be observed in A/A carriers and not G/ carriers.

## 2. Methods

### 2.1 Participants

Participants (*n* = 89, mean age = 21.85, *SD* = 1.70 females = 65) were undergraduates compensated with course credit. All participants were ethnic Chinese. The study was approved by the Psychology Programme Ethics Committee at the Nanyang Technological University and the research conducted in this study was performed in accordance with the guidelines set forth by this ethics committee (IRB Number: IRB-2015-08-020-01). Written informed consent was obtained from participants prior to the commencement of the study. All data are available at this URL: https://doi.org/10.21979/N9/IWGQ1M

### 2.2 Procedure

Participants were assessed individually where they were asked to complete a demographics questionnaire at the beginning of the study to obtain information regarding their early experiences with own- or other-race nannies. Participants were asked for the following details: (i) age at the time of the nanny experience, (ii) duration of the nanny experience and (iii) nationality of their nanny. 53 participants experienced the care of a nanny, with the beginning of receiving nanny care ranging from at birth to 10 years old (mean = 3 years old). The duration of nanny caregiving received ranged from 1 to 17.5 years (mean = 7 years). 14 participants had own-race nannies, of which 6 were a relative, while the rest of the participants had other-race nannies who were either (i) Indonesian (14) or (ii) Filipino (10). 15 participants had more than one nanny who were either (i) Indonesian and Filipino (8) or (ii) other nationalities (7).

Upon completion of the questionnaire, participants completed a visual categorization task to measure their performance in categorizing faces by race. The task was conducted using a Microsoft Surface Pro tablet with a touch screen, using E-prime 2.0 Professional (Psychology Software Tools, Sharpsburg, PA). Participants were instructed to place their hands at the side of the tablet to standardize the time needed to move their thumb to the response keys. Participants were also reminded to read the instructions carefully and perform the task as fast and as accurately as possible.

### 2.3 Experimental task

#### 2.3.1 Face stimuli

To assess participants’ categorizing ability in identifying the race of face stimuli, 4 unique Chinese faces and 4 unique Javanese faces were used as the face stimuli. Indonesian Javanese faces were used to represent other-race nannies in Singapore as Indonesians made up the majority of foreign domestic workers in Singapore - about 50.4% of foreign domestic workers (Devaraj, 2020). All faces, displayed with neutral expressions, were in colour and taken from a frontal view presented on a white background. The face images were standardized at 480 pixels (17cm) wide and 600 pixels (21cm) high with a resolution of 72 pixels per inch and were processed to be the same elliptical shape and size, with eyes and nose centrally aligned.

#### 2.3.2 Trials

The visual categorization task consisted of eight test trials where either a own- (Chinese) or other- (Javanese) race face stimulus will be presented in the middle of the display screen with two race labels. These race labels were programmed at the bottom-left or bottom-right corners of the screen as response keys. A new face was presented following the participant’s response. The faces were presented in a randomized order. To control for and minimize the effect of side preference, the position of race labels was counterbalanced such that the race labels alternated between left and right positions for each race. For instance, half of the own-race face stimuli were programmed with other-race label on the left and own-race label on the right, and vice versa. The same was programmed for Javanese face stimuli. Participants were not provided with feedback indicating whether their response was correct during the test trials.

For each stimulus, the Reaction Time (RT), i.e., the time interval between the presentation of the face and the tapping on the selected label, was measured. Then, since no differences were found between the RT of the two races (Kruskal-Wallis test: H=0.67, p=0.412), the categorization Reaction Time (ORT) was computed for each subject as the average RT across all trials.

### 2.4 OXTr genotyping

Buccal mucosa samples were collected from each participant and genotyped by a laboratory. DNA extraction was performed for each participant using the Oragene DNA purifying reagent and concentrations were assessed using spectroscopy (NanoDrop Technologies, USA). Polymerase Chain Reaction (PCR) was conducted to amplify the target OXTr gene region rs53576 using 1.5 ll of genomic DNA from the test sample, PCR buffer, 1 mM each of the forward and reverse primers, 10 mM deoxyribonucleotides, KapaTaq polymerase, and 50 mM MgCl2. The forward and reverse primers used were 5-GCC CAC CAT GCT CTC CAC ATC-3 and 5-GCT GGA CTC AGG AGG AAT AGG GAC-3. The PCR process involved (i) 15 minutes of denaturation at 95 degrees Celsius, (ii) 35 cycles at 94 degrees Celsius (30 seconds), 60 degrees Celsius (60 seconds), 72 degree Celsius (60 seconds) and 10 minutes of protraction at 72 degrees Celsius.

The division of participants into the rs53576 genotypes was: 46 (51.7%) A/A carriers, 30 (33.7%) A/G carriers, and 13 (14.6%) G/G carriers. No significant difference (X^2^(N=89, 2)=4.52, p=0.104) was observed between the distribution of participants of this study and the reference East-Asian population (A/A: 42.1%, A/G: 45.8%, G/G: 12.1% (Yates et al, 2020)).

Previous studies investigating rs53576 genotypes show a higher prevalence of G/G homozygotes particularly in European-American samples where there tends to be a skewed distribution of A/A, A/G and G/G genotypes corresponding to 12%, 38% and 50% respectively (Kim et al, 2010). Due to this skewed distribution, previous publications have largely clustered the genotype variants into G/G vs A/ carriers. However, allele frequencies and linkage disequilibrium patterns differ between Asians and Caucasians (Tost et al, 2011) and multiple studies have found that A allele frequency of rs53576 is higher in Asian populations (Luo & Han, 2014; Butovskaya et al, 2016).

Participants were therefore classified into (i) “G”: G carriers– those with at least one G allele (G/G homozygotes or A/G)) or (ii) “AA”: A/A homozygotes. The averaged distribution of the different genotypes in the Asiatic population is 45-65% for A/A homozygotes and 35-55% for G-carriers.

### 2.5 Experience with nanny

Participants were also assigned to three groups, based on the reported experience with nannies: (i) Nil group (N=36): participants with no caregiving experience; (ii) Own-race group (N=14): participants who experienced caregiving by a own-race nanny; (iii) Other-race group (N=39): participants who experienced caregiving by an other-race nanny.

A score was assigned to each group to account for the different level of experience with nannies: Nil was assigned an Experience score of 0, Own-race was assigned a score of 1 and Other-race was assigned a score of 2. The distribution of the participants between the different Experience scores is comparable in the two genetic groups: X^2^(2, N= 89) = 0.527, p=.768 (see Table 1).

**Table 1:** Distribution of the samples in the two genetic groups and types of early Nanny experience.

|  | Experience<br>(Nanny Experience) | N<br>(% of genetic group) | ORT<br>Mean<br>(SD) |
| --- | --- | --- | --- |
| AA<br>(N=46) | 0<br>(Nil) | 19<br>(41.3) | 1217.33<br>(254.76) |
|  | 1<br>(Own-race) | 6<br>(13.0) | 1043.21<br>(145.96) |
|  | 2<br>(Other-race) | 21<br>(45.7) | 958.72<br>(138.07) |
| G<br>(N = 43) | 0<br>(Nil) | 17<br>(39.5) | 1065.05<br>(150.23) |
|  | 1<br>(Own-race) | 8<br>(18.6) | 1067.42<br>(217.94) |
|  | 2<br>(Other-race) | 18<br>(41.9) | 1032.52<br>(177.96) |

### 2.6 Analytic plan

First we verified whether the two genetic groups performed differently in terms of categorization reaction times using a Kruskal-Wallis H test. Similarly, we investigated the correlation between exposure levels and categorization reaction times using a Spearman’s Rank-Order Correlation test.

A General Linear Model was fit to investigate the effect of the exposure, genetic groups and their interaction on the categorization reaction times. Post-hoc comparisons focused on separately investigating the correlation between exposure levels and categorization reaction times for the two genetic groups, using a Spearman’s Rank-Order Correlation test. Statistical significance was corrected with the Bonferroni correction method to account for multiple tests.

## 3. Results

In the preliminary analysis we found no significant difference between the two OXTR rs53576 genetic groups (H=0.04, p = 0.837, Figure 2A). A significant correlation was found between the Nanny Experience and the categorization reaction times (Spearman’ r=−0.35, p<0.001, Figure 2B).

**Figure 2.**
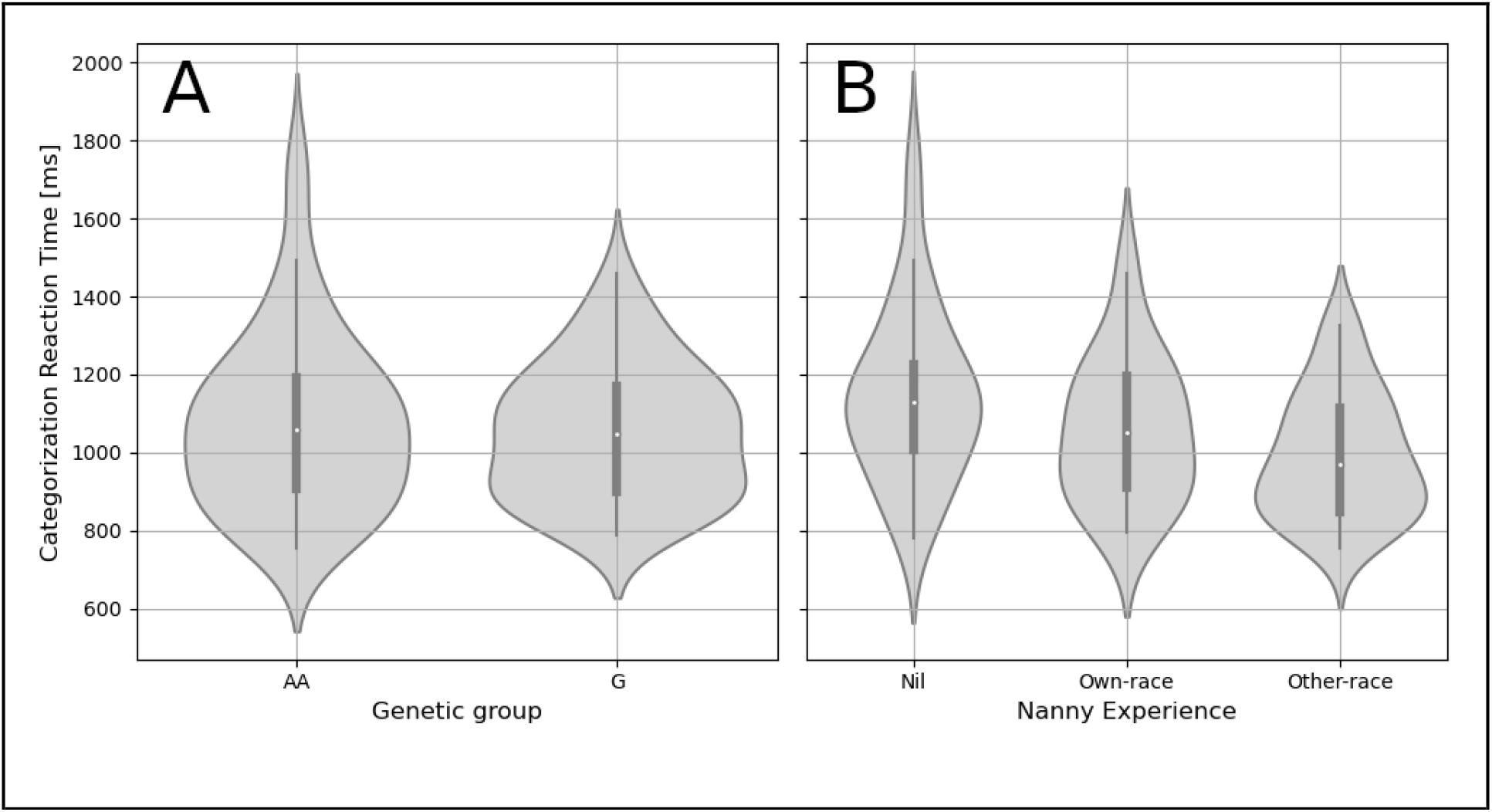
Distribution of the categorization reaction times by A. genetic group and B. Nanny Experience.

**Figure 3:**
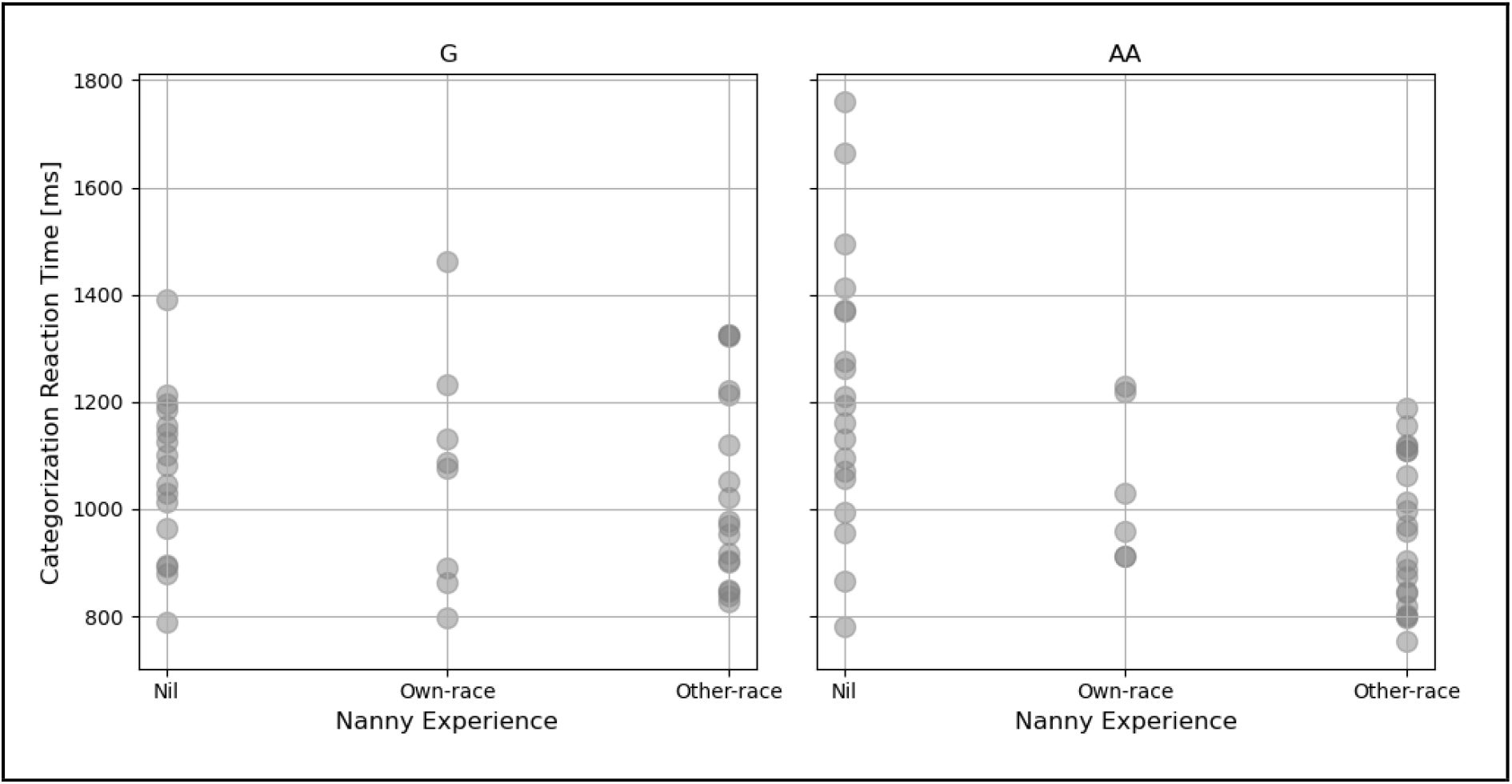
Categorization Reaction Time (ms) by different levels of exposure for the two OXTR genetic groups.

The GLM (Table 2) revealed a main effect of exposure and an interaction between exposure and genetic group. The main effect of the genetic group was not significant after the Bonferroni correction.

**Table 2:** Results of the General Linear Model to investigate the effects of exposure and OXTr genetic groups and their interaction on the categorization reaction times.

|  | Coeff | [0.025 | 0.975] | SE | z | p-value |
| --- | --- | --- | --- | --- | --- | --- |
| Intercept | 1211.18 | 1131.01 | 1291.34 | 40.9 | 29.613 | <0.001 |
| GeneticGroup | -142.56 | -258.31 | -26.804 | 59.06 | -2.414 | 0.016 |
| Experience | -129.01 | -186.32 | -71.702 | 29.24 | -4.412 | <0.001 |
| Experience:GeneticGroup | 112.648 | 28.788 | 196.508 | 42.786 | 2.633 | 0.008 |

In the post-hoc analysis we separately investigated the correlation between categorization reaction times and exposure for the two OXTR genetic groups. A significant correlation was found for the AA group (Spearman’ r=−0.52, p<0.001), but not for the G group (Spearman’ r= −0.10, p=0.533).

Participants in the AA group with Other-race nanny experience were the fastest (categorization reaction time: Mean=958 ms, SD=138 ms), while those with no nanny experience were the slowest (categorization reaction time: Mean=1217 ms, SD=254 ms) and the difference between the two groups was significant (H=11.19, p=.001).

The categorization reaction times in the G group was comparable across all groups (Nil: Mean=1065 ms, SD=150 ms; Own-race: Mean=1067 ms, SD=218 ms; Other-race: Mean=1033 ms, SD=178 ms). The difference between participants with no nanny experience and participants with Other-race nanny experience was not significant (H=0.39, p=.531).

## 4. Discussion

The present study aimed to investigate the role of both genetic and environmental factors on other-race face recognition where we examined the rs53576 OXTr genotype and early caregiving experience in Singaporean adults. Here, we proposed the inclusion of a genetics perspective to be considered in addition to existing models and theories, in investigating other-race face recognition.

First, a significant correlation between the type of nanny experience and categorisation reaction time was found where individuals with other-race nanny experience had the fastest categorisation reaction times. This finding is in line with previous findings in Rennels et al (2017) and Tham, Woo & Bremner (2018) that the face processing system can be flexible to changes in caregiving experiences in an infant’s natural environment. Notably, Tham et al (2018) found some evidence indicating that Chinese infants with additional other-race (Malay; a high frequency other-race in Malaysia) caregivers showed better recognition for other-race faces than infants without additional other-race caregivers in Malaysia. In terms of the frequency of Javanese as an other-race in Singapore, Singapore has a relatively high non-resident population in Singapore. As of 2020, there were 1.64 million non-residents, making up close to 30% of the total population of 5.69 million (Department of Statistics, 2020). Of the non-resident population, there are 252,600 foreign domestic workers, making up 18.7% of the non-resident population (Ministry of Manpower, 2020) where it has been reported that one in five Singaporean households hires a foreign domestic worker (Awang & Wong, 2019). Hence, Javanese can be considered a relatively high frequency other-race in Singapore. In addition, Javanese faces look relatively similar to Malay faces, which is the second largest racial group in Singapore. As such, in the context of the relatively large population of Javanese foreign domestic workers in Singapore, an individual needs to be able to individuate and recognise their own other-race nanny apart from other Javanese faces, leading to greater perceptual expertise derived from early experience with individuation and categorisation of Javanese faces. Since recognition and categorisation appear to share a common underlying mechanism, early experience with individuating an other-race face could result in more efficient cognitive processes involved in face categorisation and consequently, faster reaction times for race-based categorisation.

Second, although there was no significant difference in categorisation reaction times between rs53576 OXTr genotypes, a significant interaction effect of rs53576 with early experience with nannies was found. A possible explanation for this finding is the difference in prosociality between A/A homozygotes and G carriers. As discussed earlier, the G allele has previously been associated with adaptive prosocial traits. Hence, it is possible on average, G carriers have greater interracial contact with others and derive perceptual expertise from their social interactions as opposed to early caregiving experience compared to A/A homozygotes. As such, early caregiving experience constitutes a more significant source of interracial contact and exposure to other-race faces for A/A homozygotes and forms a significant avenue through which A/A homozygotes derive perceptual expertise in face categorisation. These findings suggest that the environment – early caregiving experience – may be able to compensate for a profile of less adaptive social traits and characteristics that have previously been associated with A/A carriers of rs53576. Overall, these findings also strongly suggest the importance of applying a genetics perspective to examining other-race face categorisation given the significant gene-environment interaction discovered in the present study (see Esposito, Setoh, Shinohara & Bornstein, 2017).

The findings of the present study in a multiracial societal context suggest that interracial contact and early childhood experience with other-race faces appear to influence the other-race face categorization and future studies leveraging on the nature of multiracial societies can help to provide greater insight in investigating this phenomenon. Extending from this role of early childhood experience with other-race faces, another implication of these findings could be an important consideration during child development, especially in light of findings that racial categorization of faces has been associated with implicit racial bias in children (Setoh et al, 2019) and that training children can reduce implicit racial bias (Xiao et al, 2015).

However, there are a number of limitations to the study. Firstly, early caregiving experience with a nanny was simply measured in terms of absence of nanny or presence of an other- or own-race nanny. A number of other factors such as time spent with the nanny and the relationship could possibly influence the experience that individuals would have. Hence, future studies can investigate these factors to better understand how these factors may moderate the effect of early caregiving experience on race-based face categorisation.

Secondly, early experience with nanny was measured retrospectively by asking adult participants to recall their childhood experiences. Future studies can seek to further strengthen current findings by employing a longitudinal design examining race-based face categorization in children as they develop, where the possible explanations that we proposed for our findings can be tested in terms of the relationship between the ability to individuate one’s other-race nanny and the categorisation response. These studies will also contribute to a more accurate depiction of the duration and timing of exposure to other-race nannies that must elapse for significant differences in categorisation responses with individuals without caregiving experience to arise.

## 5. Conclusions

The present study found that early nanny experience affected speed of categorisation response times, suggesting that perceptual expertise could play a role in face categorisation abilities in adults of a multiracial society. There was also preliminary evidence pointing towards gene-environment interactions influencing speed of face categorisation, specifically the rs53576 OXTr genotype. Such gene-environment interactions suggest that environmental factors could help to compensate for genetic predispositions relating to face categorisation. Findings from the present study suggest that studies on face categorisation need to take into account the demographic of the population individuals are exposed to as well as the contributing role of genetics, especially given the stark differences in both these factors in societies across the world.

## Acknowledgements

This research was supported by NAP SUG 2015 (GE), Singapore Ministry of Education ACR Tier 1 (GE; RG149/16 and RT10/19), and Singapore Ministry of Education Social Science Research Thematic Grant (MOE2016-SSRTG-017, PS).

## Author contributions

P.S., and G.E. conceptualized the study. P.S. collected the data; M.T. and J.N.F. analyzed the genetic data; A.B., analyzed the data. M.N. wrote the first draft; all the authors reviewed and edited the submitted version of the article.

## References

Anzures, G., Quinn, P. C., Pascalis, O., Slater, A. M., & Lee, K. (2010). Categorization, categorical perception, and asymmetry in infants’ representation of face race. Developmental Science, 13, 553–564.

Awang, N. & Wong, P. T. (2019, November 4). The big read: as maids become a necessity for many families, festering societal issues could come to the fore. Channel News Asia. Retrieved from https://www.channelnewsasia.com/news/singapore/maids-foreign-domestic-workers-singapore-necessity-families-12059068

Butovskaya, P. R., Lazebny, O. E., Sukhodolskaya, E. M., Vasiliev, V. A., Dronova, D. A., Fedenok, J. N., … Butovskaya, M. L. (2016). Polymorphisms of two loci at the oxytocin receptor gene in populations of Africa, Asia and South Europe. BMC Genetics, 17(17).

Department of Statistics. (2020). Population trends, 2020 (ISSN 2591-8028). Singapore: Ministry of Trade and Industry

Devaraj, S. (2020, December 17). New Indonesian maids to cost up to $3,000 more for employers in Singapore. The Straits Times. Retrieved from https://www.straitstimes.com/singapore/new-indonesian-maids-to-cost-up-to-3000-more-for-employers-in-singapore

Donaldson, Z.R. & Young, L.J. (2008). Oxytocin, vasopressin, and the neurogenetics of sociality. Science, 322: 900–904.

Dunham, Y., Chen, E. E., & Banaji, M. R. (2013). Two signatures of implicit intergroup attitudes developmental invariance and early enculturation. Psychological Science, 24(6), 860–868.

Dunham, Y., Stepanova, E. V., Dotsch, R., & Todorov, A. (2015). The development of race-based perceptual categorization: Skin color dominates early category judgments. Developmental Science, 18(3), 469–483.

Ebstein, R.P., Israel, S., Chew, S.H., Zhong, S. & Knafo, A. (2010). Genetics of human social behavior. Neuron, 65:831–844.

Esposito, G., Setoh, P., Shinohara, K., Bornstein, M. H. (2017). The development of attachment: Integrating genes, brain, behavior, and environment. Behavioral Brain Research, 325(B), 87–89.

Feng, L., Liu, J., Wang, Z., Li, J., Li, L., Ge, L., … Kang, L. (2011). The other face of the other-race effect: An fMRI investigation of the other-race face categorization advantage. Neuropsychologia, 49, 3739–3749.

Gillath, O., Shaver, P.R., Baek, J.M. & Chun, D.S. (2008). Genetic correlates of adult attachment style. Personality and Social Psychology Bulletin, 34:1396–1405.

Hugenberg, K., Young, S. G., Bernstein, M. J., & Sacco, D. F. (2010). The categorization-individuation model: An integrative account of the other-race recognition deficit. Psychological Review, 117(4), 1168–1187.

Kogan, A., Saslow, L.R., Impett, E.A., Oveis, C., Keltner, D. & Saturn S.R. (2011). Thin-slicing study of the oxytocin receptor (OXTR) gene and the evaluation and expression of the prosocial disposition. Proceedings of the National Academy of Science USA, 108: 19189–19192.

Lee, K., Quinn, P. C., Pascalis, O., & Slater, A. (2011). Development of face processing abilities. In P. D. Zelazo (Ed.), Oxford handbook of developmental psychology (Vol. 2, pp. 338–370). New York, NY: Oxford University Press.

Levin, D. T. (1996). Classifying faces by race: The structure of face categories. Journal of Experimental Psychology: Learning, Memory, and Cognition, 22, 1364–1382.

Levin, D. T. (2000). Race as a visual feature: Using visual search and perceptual discrimination tasks to understand face categories and the cross-race recognition deficit. Journal of Experimental Psychology: General, 129, 559–574.

Luo, S. & Han, S. (2014). The association between an oxytocin receptor gene polymorphism and cultural orientations. Culture and Brain, 2, 89–107.

Maddox, K. B. (2004). Perspectives on racial phenotypicality bias. Personality and Social Psychology Review, 8, 383–401.

Meyer-Lindenberg, A., Domes, G., Kirsch, P. & Heinrichs, M. (2011). Oxytocin and vasopressin in the human brain: social neuropeptides for translational medicine. Nature Reviews Neuroscience, 12: 524–538.

Ministry of Manpower. (2020). Foreign workforce numbers. Retrieved from http://www.mom.gov.sg/documents-and-publications/foreign-workforce-numbers

Ohman, A. & Mineka, S., (2001). Fears, phobias, and preparedness. Toward an evolved module of fear and fear learning. Psychol. Rev. 108 483–522.

Pascalis, O., Loevenbruck, H., Quinn, P. C., Kandel, S., Tanaka, J. W., & Lee, K. (2014). On the links among face processing, language processing, and narrowing during development. Child Development Perspective, 8(2), 65–70.

Rennels, J. L., Juvrud, J., Kayl, A. J., Asperholm, M., Gredeback, G., & Herlitz, A. (2017). Caregiving experience and its relation to perceptual narrowing of face gender. Developmental Psychology, 53, 1437–1446.

Rodin, M. J. (1987). Who is memorable to whom: A study of cognitive disregard. Social Cognition, 5(2), 144–165.

Rutland, A., Cameron, L., Milne, A., & McGeorge, P. (2005). Social norms and self-presentation: Children’s implicit and explicit intergroup attitudes. Child Development, 76(2), 451–466.

Scherer, K.R., 2001. Appraisal considered as a process of multilevel sequential checking. In: Scherer, K.R., Schorr, A., Johnstone, T. (Eds.), Appraisal Processes in Emotion:Theory, Methods, Research. Oxford University Press, New York, pp. 92–120.

Setoh, P., Lee, K. J., Zhang, L., Qian, M. K., Quinn, P. C., Heyman, G. D., & Lee, K. (2019). Racial categorization predicts implicit racial bias in preschool children. Child Development, 90(1), 162–179.

Skuse, D. H., & Gallagher, L. (2011). Genetic influences on social cognition. Pediatric research, 69(8), 85–91.

Sporer, S.L. (2001). Recognizing faces of other ethnic groups: An integration of theories. Psychology, Public Policy, and Law, 7, 36–97.

Stepanova, E. V., & Strube, M. J. (2012). The role of skin color and facial physiognomy in racial categorization: Moderation by implicit racial attitudes. Journal of Experimental Social Psychology, 48(4), 867–878.

Strategy Group. (2019). Population in Brief 2019. Republic of Singapore: Prime Minister’s Office

Tham, D. S. Y., Woo, P. J., & Bremner, J. G. (2018). Development of the other-race effect in Malaysian-Chinese infants. Developmental psychobiology, 61(1), 107–115.

Tost, H. et al. (2010). A common allele in the oxytocin receptor gene (oxtr) impacts prosocial temperament and human hypothalamic-limbic structure and function. Proceedings of the National Academy of Science, 107, 13936–13941.

Tost, H., Kolachana, B., Verchinski, B. A., Bilek, E., Goldman, A., Mattay, V., … Meyer-Lindenberg, A. (2011). Neurogenetic effects of oxtr rs2254298 in the extended limbic system of healthy caucasian adults. Biological Psychiatry, 70, e37–e39

Valentine, T., & Endo, M. (1992). Towards an exemplar model of face processing: The effects of race and distinctiveness. Quarterly Journal of Experimental Psychology A: Human Experimental Psychology, 44, 671–703.

Woo, P. J., Quinn, P. C., Méary, D., Lee, K., & Pascalis, O. (2020). A developmental investigation of the other-race categorization advantage in a multiracial population: Contrasting social categorization and perceptual expertise accounts. Journal of experimental child psychology, 197, 104870.

Xiao, W. S., Fu, G., Quinn, P. C., Qin, J., Tanaka, J., Pascalis, O., & Lee, K. (2015). Individuation training with other-race faces reduces preschoolers’ implicit racial bias: A link between perceptual and social representation of faces in children. Developmental Science, 18(4), 655–663.

Yates, A. D., Achuthan, P., Akanni, W., Allen, J., Allen, J., Alvarez-Jarreta, J., … & Bhai, J. (2020). Ensembl 2020. Nucleic acids research, 48(D1), D682–D688.

Zhao, L., & Bentin, S. (2008). Own- and other-race categorization of faces by race, gender, and age. Psychonomic Bulletin & Review, 15, 1093–1099.

